# Functional inactivation of pulmonary MAIT cells following 5-OP-RU treatment of non-human primates

**DOI:** 10.1101/2021.01.29.428844

**Authors:** Shunsuke Sakai, Nickiana E. Lora, Keith D. Kauffman, Danielle E. Dorosky, Sangmi Oh, Sivaranjani Namasivayam, Felipe Gomez, Joel D. Fleegle, Tuberculosis Imaging Program, Cecilia S. Lindestam Arlehamn, Alessandro Sette, Alan Sher, Gordon J. Freeman, Laura E. Via, Clifton E. Barry, Daniel L. Barber

## Abstract

Targeting MAIT cells holds promise for the treatment of different diseases and infections. We previously showed that treatment of *Mycobacterium tuberculosis* infected mice with 5-OP-RU, a major antigen for MAIT cells, expands MAIT cells and enhances bacterial control. Here we treated *M. tuberculosis* infected rhesus macaques with 5-OP-RU intratracheally but found no clinical or microbiological benefit in *M. tuberculosis* infected macaques. In fact, after 5-OP-RU treatment MAIT cells did not expand, but rather upregulated PD-1 and lost the ability to produce multiple cytokines, a phenotype resembling T cell exhaustion. Furthermore, we show that vaccination of uninfected macaques with 5-OP-RU+CpG instillation into the lungs also drives MAIT cell dysfunction, and PD-1 blockade during vaccination partly prevents the loss of MAIT cell function without facilitating their expansion. Thus, in rhesus macaques MAIT cells are prone to the loss of effector functions rather than expansion after TCR stimulation *in vivo*, representing a significant barrier to therapeutically targeting these cells.

## INTRODUCTION

Mucosal Associated Invariant T (MAIT) cells are restricted by the nonpolymorphic MHC class I-like molecule MR1 and express TCRs specific for small molecule metabolites produced by microbes (1, 2). 5-OP-RU, a derivative of intermediates produced during bacterial riboflavin biosynthesis, is major MR1 ligand and stimulatory MAIT cell antigen that is recognized by a majority of MR1-restricted T cells (3–5). MAIT cells display pro-inflammatory, cytotoxic, as well as tissue-repair properties (6–8). Given the non-polymorphic nature of MR1 there is interest in targeting MAIT cells through vaccination or as therapies for the treatment of various conditions including cancer and infections. Indeed, using mouse models, MAIT cells have been shown to be important for the control of certain bacterial infections (9–11), and they can easily be driven to expand to large numbers *in vivo* via vaccination with antigen and adjuvant (7, 12).

Targeting MAIT cells may be useful during *Mycobacterium tuberculosis* (Mtb) infection (13). Recently, it was shown that MR1 deficient mice have no defect in bacterial control or survival after Mtb infection (14, 15), and the presence of large populations of MAIT cells at the time of Mtb exposure has no impact on host resistance (14–16). Therefore, the murine model data indicate that pre-infection vaccination of MAIT cells may not be beneficial for Mtb infection. However, we showed that treatment of mice harboring a chronic Mtb infection with 5-OP-RU was able to reduce bacterial loads ~10 fold in 3 weeks in a manner dependent on IL-17A (14), indicating that post-exposure stimulation of MAIT cells may be a promising host directed therapy for tuberculosis. While there is significant overlap in the gene expression patterns of human and mouse MAIT cells (7), there are several key differences between murine and nonhuman primate (NHP)/human MAIT cells. For example, the majority of mouse MAIT cells are CD4^-^CD8^-^ IL-17A-producing MAIT17 cells, while most human and macaque MAIT cells are CD8^+^ IFN-γ-producing MAIT1 cells (8, 17–19). Moreover, mice have several limitations in their ability to model human tuberculosis (TB). In contrast, Mtb infection of macaques recapitulates most features of human TB, and macaques are considered the gold standard pre-clinical model of TB (20). Therefore, we tested the potential therapeutic efficacy of 5-OP-RU instillation into the lungs of Mtb infected rhesus macaques.

In contrast to mice, there was no therapeutic benefit observed with 5-OP-RU treatment of macaques with TB. MAIT cells in these animals failed to expand, upregulated high levels of PD-1 and became functionally impaired. To ask if the loss of function was due to Mtb infection or PD-1, we vaccinated macaques with 5-OP-RU+CpG instillation in the lungs in the presence or absence of PD-1 blocking antibody and examined MAIT cell expansion and function. Again, MAIT cells failed to expand and lost the ability to produce cytokines. PD-1 blockade during 5-OP-RU+CpG vaccination did not result in MAIT cell expansion but partly alleviated the loss of cytokine producing ability. This study provides insight into basic in vivo MAIT cell biology in NHPs and identifies a major barrier to the translatability of MAIT cell directed therapies. Unlike mice, administration of 5-OP-RU during infection or with adjuvant in macaques fails to expand MAIT cells and instead leads to MAIT cell dysfunction. Strategies to drive MAIT cell expansion in macaques are needed to test their therapeutic potential during TB and other diseases.

## RESULTS

### 5-OP-RU treatment in Mtb-infected macaques does not enhance control of the infection

To examine the therapeutic efficacy of MAIT cell stimulation during Mtb infection, macaques were treated intratracheally with PBS or 5-OP-RU from week 6 to week 14 post-infection (Figure 1A). Necropsies were planned for week 15-16. All five PBS treated control animals survived until the pre-determined endpoint, however, three of the five 5-OP-RU treated animals developed acute signs of disease (cough and labored breathing) and were humanly euthanized early (Figure 1B). All animals in both groups had relatively stable lung disease as measured by [^18^F]-FDG-PET/CT imaging of the lungs (Figure 1C and D). Upon necropsy, bacterial loads in individually resected granulomas were measured. Granulomas from PBS and 5-OP-RU treated macaques had similar overall numbers of bacteria (Figure 1E). However, when control animals were compared to just the two 5-OP-RU treated macaques which continued until the week15/16 pre-determined endpoint, there was a slight decrease in the numbers of bacteria in treated animals (Figure 1E). Two of the 5-OP-RU treated macaques that were euthanized early (DHLR at week 7 and DHAJ at week 10) displayed slightly increased bacterial loads in the granulomas, but we could not distinguish whether the increase in the number of bacteria was due to the earlier necropsy or 5-OP-RU treatment (Figure 1E). Moreover, there was no significant difference in the number of bacteria in the spleens of both groups (Figure 1F).

**Figure 1.**
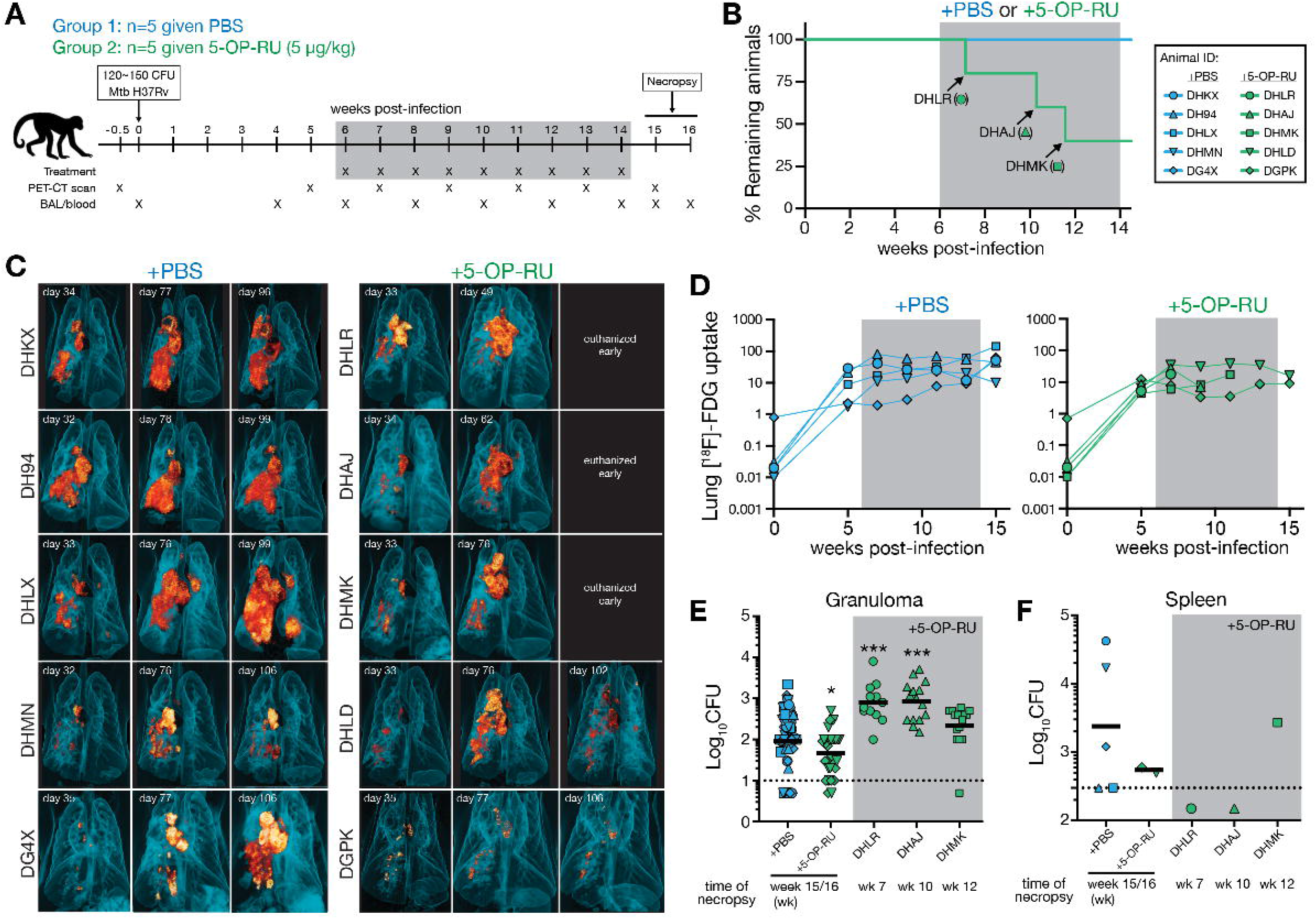
5-OP-RU treatment in Mtb-infected macaques does not enhance control of the infection. (**A**) Ten animals were infected with 120-150 colony forming units (CFU) of Mtb H37Rv. Starting 6 weeks post-infection, animals were intratracheally treated with PBS (n=5) or 5-OP-RU (n=5). Animals were necropsied at weeks 15 or 16 post-infection. (**B**) Percentage of remaining animals in the 2 groups were plotted against the weeks after Mtb infection. Three 5-OP-RU treated animals (DHLR, DHAJ, DHMK) were humanly euthanized early due to the development of severe pulmonary distress. (**C**) Three-dimensional volume renderings of PET/CT scans of each animal at week 5 and 11 post-infection and at necropsy. (**D**) The lung total [^18^F]-FDG uptake (total lesion glycolysis, TLG) value in PBS *(left)* and 5-OP-RU treated *(right)* animals. Bacterial loads in the granulomas (**E**) and spleens (**F**) shown in pooled animals by experimental group and individual animals euthanized early. Each symbol represents an individual tissue sample from indicated animal. Dotted line indicates limit of detection. \**p* < 0.05, \*\*\**p* < 0.001.

Since 3 of the 5-OP-RU treated macaques (DHLR, DHAJ, DHMK) had acute clinical signs after treatment, we assessed the levels of airway constriction in the animals by measuring the maximum diameter of bronchi on CT images collected at baseline and during Mtb infection. The three 5-OP-RU treated animals that were euthanized early showed rapid reductions in airway diameter after 5-OP-RU treatment relative to most of the PBS control treated animals and other 5-OP-RU treated 2 animals that did not develop severe signs of distress (Figure 2A and B, S2). The bronchoconstriction was not associated with either higher bacterial loads in the pulmonary lymph nodes (LNs) (Figure 2C) or overall LN [^18^F]-FDG uptake on PET/CT scans (Figure 1C and 2D). However, a trend was observed in which animals with bronchoconstriction had at least one very cellular pulmonary LN at necropsy relative to the 5-OP-RU treated animals that did not develop respiratory distress or the PBS treated animals (Figure 2E). On visual inspection at necropsy, it was apparent that the enlarged LNs were impinging on the bronchus. Therefore, it seems likely that lymphadenopathy-associated airway constriction and not expansive tubercular lung lesions was responsible for the acute clinical signs that led to the early euthanasia of these three macaques treated with 5-OP-RU. Rhesus macaques have been shown previously to be particularly susceptible to lethal LN pathology (21, 22), so it is not clear if the 5-OP-RU treatment was the direct cause of the bronchoconstriction. Regardless, 5-OP-RU treatment during Mtb infection did not have a clinical or microbiological benefit in rhesus macaques.

**Figure 2.**
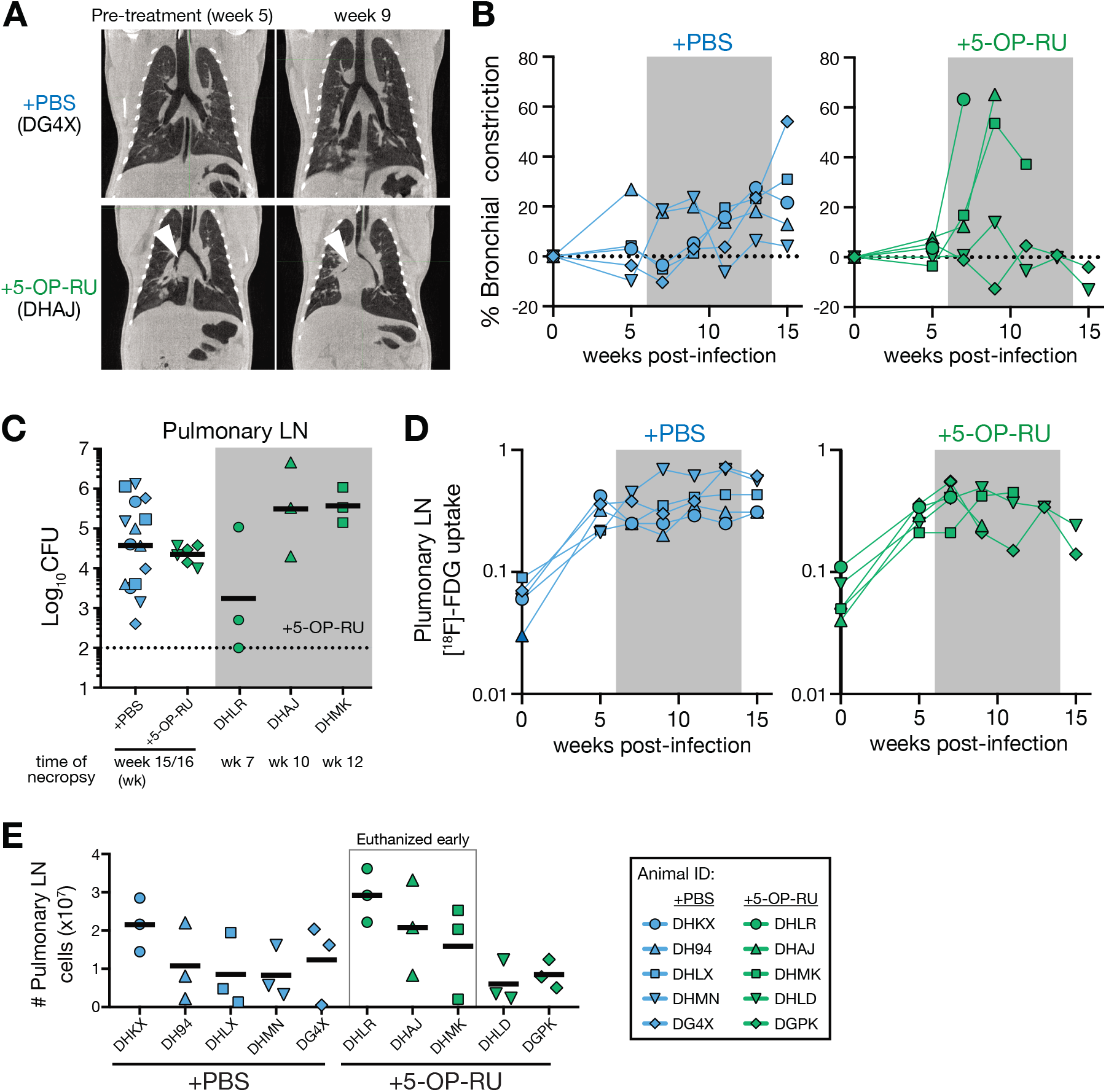
Early necropsies of 5-OP-RU treated animals were associated with bronchoconstriction. (**A**) Example chest CT images from PBS (*top*) and 5-OP-RU *(bottom)* treated animals chosen to illustrate the bronchial constriction. (**B**) Percent change of airway constriction of PBS (*left*) and 5-OP-RU treated *(right)* animals. Airway constriction was determined by comparing the diameter of bronchi at baseline and after infection. (**C**) Bacterial loads in the pulmonary LNs shown in pooled animals by experimental group and individual animals euthanized early. Each symbol represents an individual tissue sample from indicated animal. Dotted line indicates limit of detection. (**D**) The lung total [^18^F]-FDG uptake (total lesion glycolysis, TLG) value in PBS (*left*) and 5-OP-RU treated *(right)* animals. (**E**) Number of total viable cells within pulmonary LNs. Each symbol represents an individual tissue from indicated animal.

### MAIT cells become activated but do not expand after 5-OP-RU instillation

We next measured the frequency of MAIT cells in the bronchoalveolar lavage (BAL) and peripheral blood using rhesus macaque MR1/5-OP-RU tetramers. Consistent with the previous findings (23), the frequency of MAIT cells in the airway and blood did not change in the PBS treated group after Mtb infection (Figure 3A to D). In contrast to previous reports in the mouse model showing large increases in MAIT cells after 5-OP-RU administration (14, 15), 5-OP-RU treatment in the infected macaques did not result in the expansion of airway or circulating MAIT cells (Figure 3A to D). To examine whether MAIT cells entered cell cycle, we measured Ki-67 expression in MAIT cells isolated from the BAL and blood. While there was a small increase in the Ki-67 expression by MAIT cells in the BAL at 4 weeks post-infection, we did not observe clear differences in their Ki-67 expression between PBS and 5-OP-RU treated groups during treatment (Figure 3C and D). In contrast to the BAL, MAIT cells in the blood significantly upregulated Ki-67 at 4 weeks post-infection in both animal groups (Figure 3E). While the levels of Ki-67 expression in blood borne MAIT cells then decreased back toward baseline in the PBS treated group, there was a clear increase in Ki-67^+^ MAIT cells after 5-OP-RU treatment (Figure 3E).

**Figure 3.**
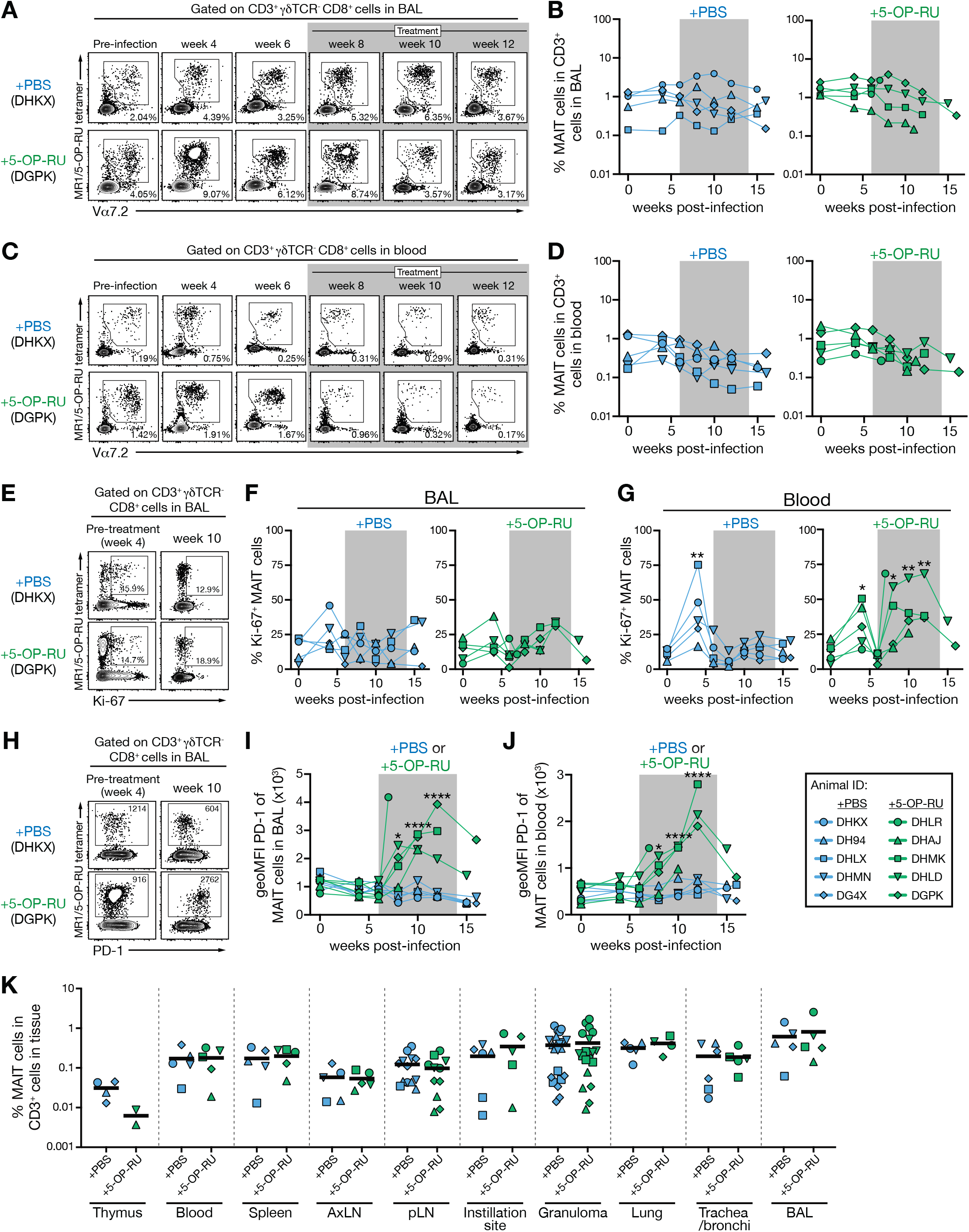
MAIT cells become activated but do not expand after 5-OP-RU instillation. (**A**) Example FACS plots of MR1/5-OP-RU tetramer staining of cells in the bronchoalveolar lavage (BAL). (**B**) Kinetics of the frequency of MAIT cells among CD3 ^+^ cells in the airway. (**C**) Example FACS plots of MR1/5-OP-RU tetramer staining of cells in the blood. (**D**) Kinetics of the frequency of MAIT cells among CD3 ^+^ cells in the blood. (**E**) Example FACS plots of Ki-67 expression on MAIT cells in the BAL from week 4 and 10 post-infection. (**F and G**) Summary graphs showing the percentage of Ki-67^+^ MAIT cells in the BAL (F) or blood (G). (**H**) Example FACS plots of PD-1 expression on MAIT cells in the BAL from week 4 and 10 post-infection. (**I and J**) Summary graphs showing the geometric mean fluorescent intensity (geoMFI) of PD-1 on MAIT cells in the BAL (I) or blood (J). (**K**) Percentage of MAIT cells in the indicated tissues at necropsy. Each symbol represents an individual tissue sample from indicated animal. \**p* < 0.05, \*\**p* < 0.01, \*\*\*\**p* < 0.0001.

PD-1 is induced by TCR signaling, so we next examined its expression to ask if MAIT cells were stimulated through their TCR after infection and 5-OP-RU treatment. While PD-1 expression by MAIT cells was not changed in the BAL or blood of control animals, 5-OP-RU treatment led to a striking upregulation of PD-1 by MAIT cells in both compartments (Figure 3H to J), indicating that MAIT cells clearly received antigen stimulation through their TCRs after 5-OP-RU treatment. At necropsy, we examined multiple tissue sites to ask if MAIT cells expanded in any tissue. There was also no impact of 5-OP-RU administration on the frequency of MAIT cells in the thymus, spleen, pulmonary LNs, granulomas or instillation site lesions (Figure 3K). Overall, these data demonstrate that 5-OP-RU treatment in Mtb infected rhesus macaques indeed activates MAIT cells via TCR stimulation but fails to induce expansion of MAIT cells in the tissues.

### 5-OP-RU treatment leads to MAIT cell dysfunction in Mtb infected macaques

We next evaluated the impact of 5-OP-RU treatment on the function of MAIT cells. Cells from the BAL or blood were stimulated with 5-OP-RU or PMA/ionomycin, and production of IFN-γ, TNF-⍰, IL-17A and GM-CSF was measured by intracellular cytokine staining (Figure 4A). The vast majority of MAIT cells in both blood and BAL produced cytokines after stimulation with PMA/ionomycin (Figure 4B to D). However, prior to instillation of 5-OP-RU, there were major differences in the responses of BAL vs blood MAIT cells after *in vitro* 5-OP-RU stimulation. Approximately 75% of MAIT cells in the BAL produced cytokines upon 5-OP-RU stimulation while only ~10% of circulating MAIT cells were able to respond (Figure 4B to D). After the macaques were treated, BAL MAIT cells in animals receiving 5-OP-RU rapidly lost the ability to respond to *in vitro* restimulation with 5-OP-RU, while MAIT cells in PBS treated macaques maintained their ability to produce cytokines (Figure 4B to D). In contrast, PMA/ionomycin induced similar levels of cytokine production by MAIT cells in both PBS and 5-OP-RU treated macaques, although there was a trend for a reduction at very late time points (Figure 4B to D). MAIT cells from pulmonary LNs, instillation site lesions in the lungs, the BAL and individually resected granulomas isolated at necropsy also showed a reduction in cytokine producing function after *in vitro* restimulation with 5-OP-RU and to a lesser extent after restimulation with PMA/ionomycin (Figure 4E and F).

**Figure 4.**
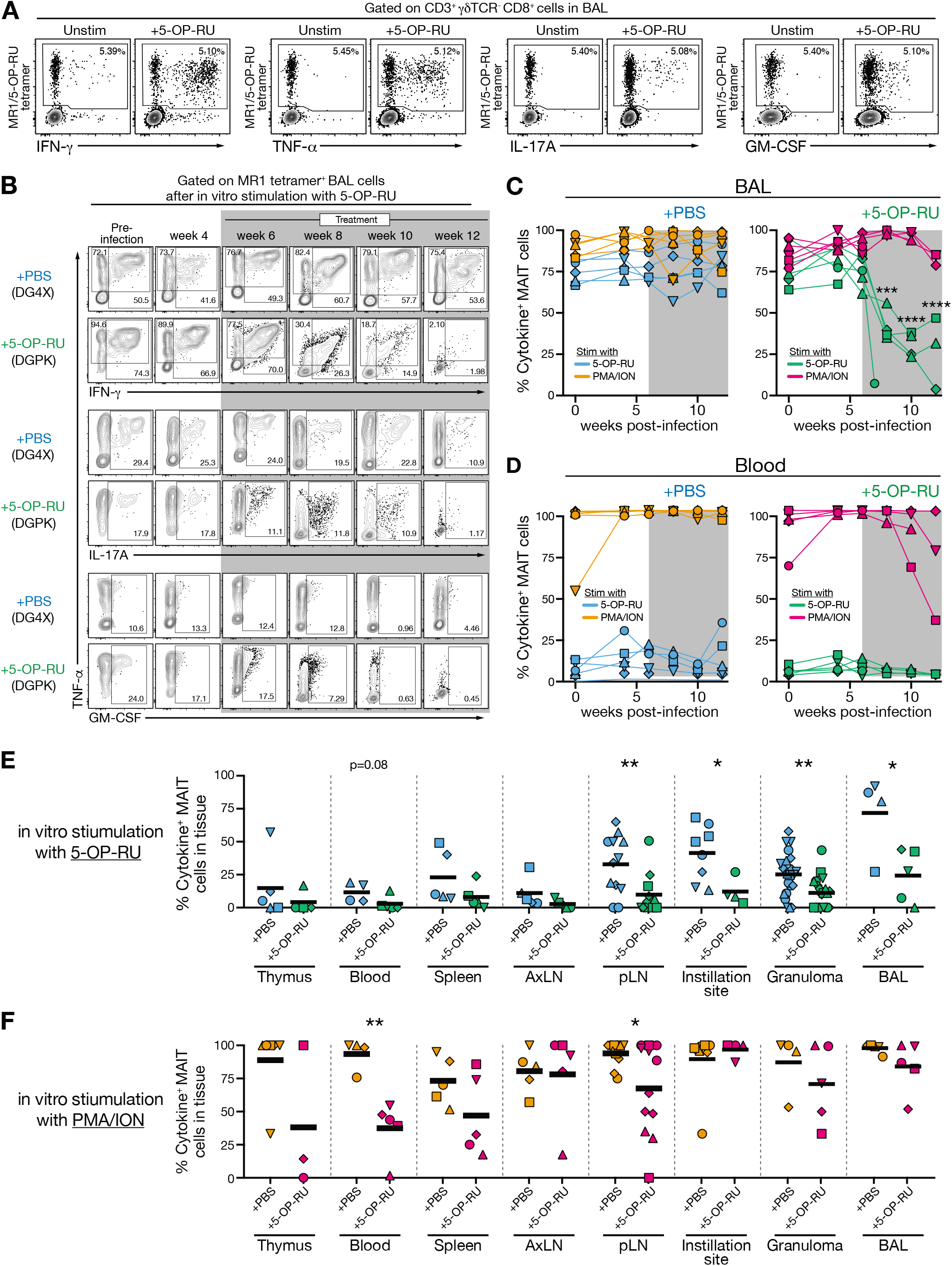
5-OP-RU treatment leads to MAIT cell dysfunction in Mtb infected macaques. (**A**) Example FACS plots of intracellular cytokine staining of MR1/5-OP-RU tetramer^+^ cells from the BAL (animal ID: DH4X) following in vitro stimulation with 5-OP-RU for 6 hours. (**B**) Example FACS plots of intracellular cytokine staining of MAIT cells from the BAL following in vitro stimulation with 5-OP-RU. (**C and D**) Summary graphs of percentage of MAIT cells that are ether IFN-γ^+^, TNF-⍰^+^, IL-17A^+^ or GM-CSF^+^ in the BAL (B) or blood (C) following in vitro stimulation with 5-OP-RU or PMA/ionomycin (PMA/ION). (**E and F**) Graphs showing the percentage of MAIT cells that are ether IFN-γ^+^, TNF-⍰^+^, IL-17A^+^ or GM-CSF^+^ following in vitro stimulation with 5-OP-RU (D) or PMA/ionomycin (E) in the indicated tissues at necropsy. Each symbol represents an individual tissue sample from indicated animal. \**p* < 0.05, \*\**p* < 0.01, \*\*\**p* < 0.001, \*\*\*\**p* < 0.0001.

These data demonstrate several important points. First, prior to infection MAIT cell function after 5-OP-RU stimulation varies substantially between different tissues, with lung MAIT cells displaying very high levels of responsiveness. Second, treatment of Mtb infected macaques with 5-OP-RU leads to a striking reduction in pulmonary MAIT cell functionality. Lastly, the functional defect can be overcome to a certain extent if the TCR is bypassed via stimulation with second messengers like PMA/ionomycin. Therefore, rather than the expected increase in MAIT cell responses after 5-OP-RU treatment, MAIT cells entered an exhaustion-like state.

### 5-OP-RU treatment of Mtb infected macaques does not enhance conventional adaptive immune responses

We next examined the impact of 5-OP-RU treatment on the conventional adaptive immune response. We found that there was no difference in the kinetics of Mtb-specific CD4 T cell response in the airways as measured by intracellular cytokine staining for IFN-γ and TNF-⍰ after restimulation with Mtb peptide megapools (Figure 5A and B). Likewise, there was no apparent impact of 5-OP-RU treatment on the kinetics of the Mtb-specific CD8 T cells in the BAL (Figure 5C and D). There was also no impact of 5-OP-RU administration on the magnitude of Mtb-specific CD4 and CD8 T cells in the blood, spleen, LNs, granulomas or instillation site lesions at necropsy (Figure 5E and F). A previous study found that MAIT cells may directly provide help to B cells in rhesus macaques (24), so we also measured Mtb-specific IgG responses in serum. However, there was no clear difference in antibody responses between treated and untreated animals (Figure 5G). Collectively, these data show that stimulation of MAIT cells during Mtb infection did not boost Mtb-specific conventional adaptive immune responses.

**Figure 5.**
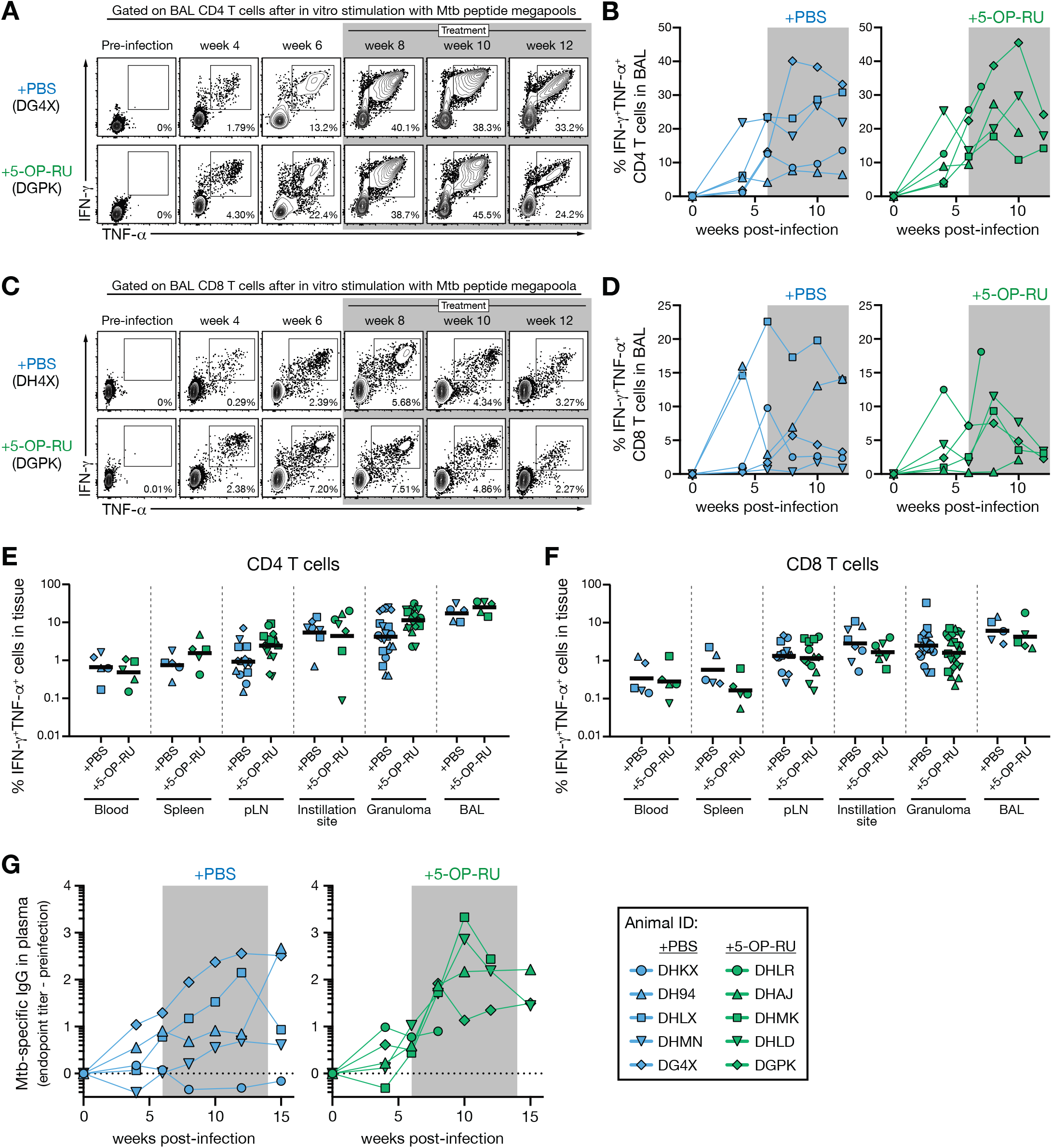
5-OP-RU treatment of Mtb infected macaques does not enhance conventional adaptive immune responses. (**A**) Example FACS plots of intracellular cytokine staining of CD4 T cells from the BAL following *in vitro* stimulation with MTB300 peptide megapool. (**B**) Summary graphs of percentage of CD4 T cells in the BAL that are IFN-γ^+^TNF-EI^+^ following in vitro stimulation. (**C**) Example FACS plots of intracellular cytokine staining of CD8 T cells from the BAL following in vitro stimulation with the MHC-I Mtb peptide megapool. (**D**) Summary graphs of percentage of CD8 T cells in the BAL that are IFN-γ^+^TNF-⍰^+^ following in vitro stimulation. (**E and F**) Graphs showing the percentage of IFN-γ^+^TNF-⍰^+^ CD4 T cells (E) or CD8 T cells (F) in the indicated tissues at necropsy. Each symbol represents an individual tissue sample from indicated animal. (**G**) Mtb-specific antibody response following infection. Plasma levels of IgG specific to Mtb antigens were calculated by subtracting background values obtained in the pre-infection samples.

### 5-OP-RU treatment does not cause major alterations in the intestinal microbiota

Due to the role of MAIT cells at mucosal sites and the ability of MAIT cells to recognize microbe-derived molecules, a close interplay between MAIT cells and the microbiota has been hypothesized (25). Indeed, recent work has demonstrated that the microbiota plays a critical role in the development of MAIT cells in mouse models (6, 26). Therefore, we sought to investigate if treatment with 5-OP-RU affects the microbiome. Fecal samples were collected from all 10 macaques prior to infection and following infection and treatment. The composition of the microbiota was characterized via 16S rRNA sequencing. We found that alpha-diversity (with-in sample diversity) of the microbiota varied over the course of the experiment in both PBS and 5-OP-RU groups (Supplemental Figure 1A, left panel). However, we did not find the alpha-diversities of the microbiome from the 5-OP-RU or PBS treated timepoints to be significantly different from each other or from their respective pre-infection and infected microbiomes (Supplemental Figure 1A, right panel). Similarly, the community structure of the microbiota following 5-OP-RU treatment was not significantly different from that of PBS treated animals (Supplemental Figure 1B). However, visualization of the relative abundance of microbial taxa over the experimental time course indicated alterations in the composition of the gut flora, albeit not consistent between animals within a treatment group (Supplemental Figure 1C). Several taxa were identified to be differently abundant with comparing the PBS or 5-OP-RU treated animals to their respective pre-infection microbiomes (Supplemental Figure 1D). Importantly, we found only three bacterial families, Gastranaerophilales, Family XIII and Paludibacteraceae to be enriched and one family, vadin BE97, to be depleted following 5-OP-RU treatment in comparison to PBS administration (Supplemental Figure 1D). Overall, these analyses reveal that the 5-OP-RU treatment results in only minimal alterations in the intestinal microbiota of rhesus macaques.

### PD-1 blockade partially restores MAIT cell function but not expansion after 5-OP-RU+CpG vaccination in uninfected macaques

In a separate experiment, we next vaccinated uninfected macaques with the same regimen of 5-OP-RU+CpG as we previously used in our murine model experiments (Figure 6A). This allowed us to test three possible explanations for the poor MAIT responses after 5-OP-RU treatment of Mtb infected macaques. First, treating uninfected macaques allowed us to ask if the Mtb infection itself inhibited the ability of MAIT cells to respond. Second, we lowered the dose tenfold to test the possibility that our previous results were due to a high zone tolerance-like effect. Lastly, MAIT cells have been shown to be regulated by PD-1 during human TB (27), so we included a group of macaques that received anti-PD-1 blocking antibody at the time of 5-OP-RU vaccination to ask if the upregulation of PD-1 was responsible for the poor MAIT cell responses *in vivo.*

**Figure 6.**
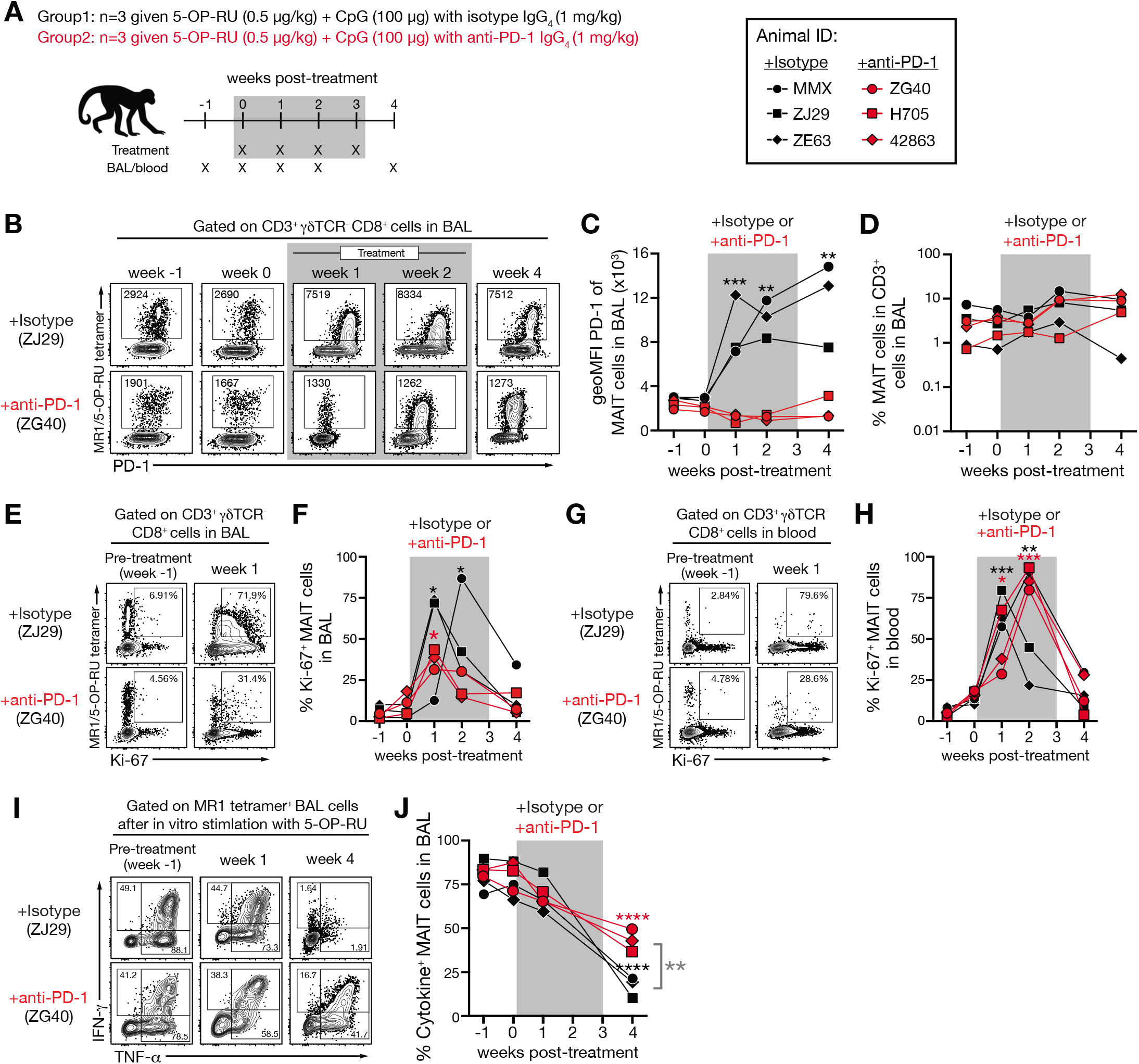
PD-1 blockade partially restores MAIT cell function but not expansion after 5-OP-RU+CpG vaccination in uninfected macaques. (**A**) Uninfected animals were treated intratracheally with 5-OP-RU+CpG along with either rhesus macaque IgG_4_ isotype control or primatized anti-PD-1 antibody. (**B**) Example FACS plots of PD-1 expression on MAIT cells in the BAL. (**C**) Summary graph showing the geoMFI of PD-1 on MAIT cells in the BAL. (**D**) Kinetics of the frequency of MAIT cells among CD3 ^+^ cells in the BAL. (**E**) Example FACS plots of Ki-67 expression on MAIT cells in the BAL from pretreatment and week 1 post-treatment. (**F**) Percentage of Ki-67^+^ MAIT cells in the BAL. (**G**) Example FACS plots of intracellular cytokine staining of MAIT cells from the BAL following in vitro stimulation with 5-OP-RU. (**H**) Percentage of MAIT cells that are ether IFN-γ^+^, TNF-⍰^+^ or IL-17A^+^ in the BAL following in vitro stimulation with 5-OP-RU. \**p* < 0.05, \*\**p* < 0.01, \*\*\**p* < 0.001, \*\*\*\**p* < 0.0001.

5-OP-RU+CpG vaccination resulted in a striking upregulation of PD-1 in isotype control treated animals indicating that MAIT cells were effectively stimulated by this vaccination (Figure 6B and C). PD-1 staining was greatly reduced on MAIT cells from animals treated with anti-PD-1, presumably as a consequence of cell surface PD-1 being occupied by the PD-1 blocking antibody (Figure 6B and C). Similar to what was observed after treatment of Mtb infected macaques, MAIT cells failed to expand in the airways after vaccination, and anti-PD-1 treatment had no effect on MAIT cell frequencies (Figure 6D). Although MAIT cells did not expand, Ki-67 was significantly upregulated on MAIT cells in both the BAL and blood after vaccination (Figure 6E to H). PD-1 blockade along with vaccination may have slightly reduced Ki-67 expression in the BAL MAIT cells and delayed its peak expression in the circulating MAIT cells, but there was relatively little difference in Ki-67 expression in the animals receiving anti-PD-1. Collectively, these data show that 5-OP-RU+CpG vaccination of rhesus macaques leads to MAIT cell recognition of the antigen and induces them to enter cell cycle but fails to result in an expansion of MAIT cells (Figure 6B and C).

At different time point after vaccination, we also restimulated cells from the airways with 5-OP-RU *in vitro* and measured MAIT cell functionality with intracellular cytokine staining. Again, we found that MAIT cells underwent a significant reduction in their cytokine producing potential after 5-OP-RU administration (Figure 6I and J). Interestingly, PD-1 blockade partially rescued the function of MAIT cells at 4 weeks post-treatment, indicating that loss of effector functions but not proliferation was in part due to PD-1 expression. Thus, the lack of MAIT cell expansion and antigen-driven reduction in MAIT cell functionality was not due to the presence of the Mtb infection and was very similar despite a 10-fold reduction in 5-OP-RU dose. Moreover, PD-1 expression was partly responsible for the loss of MAIT cell function but had no role in their poor expansion.

## DISCUSSION

We previously showed that treating Mtb infected mice with the MAIT cell TCR agonist 5-OP-RU leads to a large expansion of MAIT cells and reduction in lung bacterial loads (14), prompting us to further evaluate the translatability of MAIT cell directed therapy during TB in the rhesus macaque model. While no control animals were lost prior to the predetermined endpoint, three of five treated animals developed distress relating to bronchoconstriction after 5-OP-RU instillation and needed to be humanely euthanized early. In all cases this appeared to be due to enlarged LNs impinging on airways, so 5-OP-RU treatment may have exacerbated disease in these animals. However, rhesus macaques have been shown to be particularly prone to developing TB lymphadenopathy (21, 22), bacterial loads were not substantially increased in the animals that developed early disease and there was no obvious biological effect on MAIT cells to which we could ascribed this outcome. Therefore, we interpret this result with caution. A larger study would be needed to more definitively establish a causal link between 5-OP-RU treatment and lymphadenopathy in macaques. Regardless, it is clear 5-OP-RU treatment in macaques did not lead to the beneficial outcomes we observed in mice. 5-OP-RU treatment failed to expand MAIT cells and instead only drove them into a functionally exhausted-like state. This was not a result of the Mtb infection, as vaccination of uninfected macaques by instillation of 5-OP-RU (adjuvanted with CpG) into the lungs also did not increase the frequency of MAIT cells and likewise resulted in their loss of cytokine producing ability. Functional inactivation of MAIT cells following *in vivo* administration of 5-OP-RU during TB and vaccination was associated with upregulation of the co-inhibitory receptor PD-1. Therefore, we asked if PD-1 blockade could boost MAIT cell responses following pulmonary 5-OP-RU+CpG vaccination. While PD-1 blockade did not result in MAIT cell expansion, it did partly alleviate their loss of function. Therefore, rather than the expansion of MAIT cells normally seen in mice after administration of 5-OP-RU, in rhesus macaques MAIT cells fail to proliferate and undergo functional inactivation of their cytokine producing activity. Our study reveals an important conundrum. That is, while MAIT cells are very weakly stimulated by Mtb infection (14, 15, 23, 28), attempting to drive MAIT responses by treating macaques with additional TCR ligand only decreases their functionality.

Previous studies have documented defects in MAIT cell function during infection, cancer and sepsis (29–38). Our data showing MAIT cell functional defects directly resulting from TCR stimulation indicates that part of the MAIT dysfunction observed in these settings may result from increased MAIT antigen stimulation, rather than disease-related bystander processes. The loss of MAIT cell function that occurs in these reports and here after 5-OP-RU treatment resembles the exhaustion of virus and tumor-specific conventional CD8 T cells. However, it is unclear if this is the same process for several reasons. Exhaustion of conventional CD8 T cells is a unique differentiation state that occurs during a period of chronic antigenic stimulation following an initial clonal burst of highly functional effector cells (39). Following 5-OP-RU treatment, MAIT cells never underwent an expansion phase. They simply began to lose effector functions quickly after *in vivo* stimulation. Moreover, the ability of MAIT cells in different tissues to produce cytokine following 5-OP-RU stimulation is highly variable, confounding comparisons to exhaustion. At baseline, MAIT cells in the BAL were the most functional and those in the blood the least. Even after lung MAIT cells had undergone this functional inactivation, they still produced more cytokine than blood cells prior to treatment. In other words, it is difficult to call lung MAIT cells exhausted if their counterparts in circulation are even less functional at their best. Therefore, it is not clear if the pulmonary MAIT cells present after 5-OP-RU treatment are truly exhausted as understood for conventional T cells. In future experiments, it will be important to molecularly characterize MAIT cells before and after 5-OP-RU-treatment in vivo to identify mechanisms leading to their dysfunction and better understand the nature of this unusual T cell inactivation process in relation to bona fide exhaustion of conventional CD8 T cells.

PD-1 is a key mediator of the exhaustion of conventional peptide-specific CD8 T cells, and several biologics targeting PD-1 or its ligands are used in the treatment of various cancers. Here we saw a striking upregulation of PD-1 on MAIT cells in the lungs as well as circulation after 5-OP-RU treatment. PD-1 blockade during 5-OP-RU+CpG vaccination did at least partly prevent the reduction in MAIT cell function. Therefore, while PD-1 does have a role in MAIT dysfunction, it seems likely that other regulatory pathways also have significant contributions to the downregulation of MAIT cell effector functions after in vivo stimulation. Interestingly, the defect in cytokine production after 5-OP-RU treatment was only observed when the cells were restimulated with 5-OP-RU, and the cells displayed equivalent higher cytokine production as controls upon PMA/ionomycin restimulation, indicating that the defect relates to signaling through the TCR. It is possible that multiple co-inhibitory receptors cooperate in MAIT cell inactivation, as MAIT cells have been shown to express several different immune checkpoint molecules (31, 33, 34, 40, 41).

Our original goal was to drive large populations of MAIT cells in the lungs of macaques and examine the impact on Mtb infection. However, MAIT cells failed to increase in number after instillation of 5-OP-RU, and PD-1 blockade during antigenic stimulation did not lead to MAIT cell expansion. In mice, MAIT cells can readily be driven to expand ~100 fold, as long as 5-OP-RU is properly adjuvanted (7, 12, 14–16). However, unlike mice, it is not clear that NHP or human MAIT cells are able to undergo similar dramatic expansions *in vivo.* Studies of human vaccination with *Shigella dysenteriae 1* (42) or BCG (18), as well as controlled human bacterial infection studies with *Salmonella enterica* serovar Typhi (43) and *Salmonella enterica* serovar Paratyphi A (44) have consistently found that MAIT cells become activated but do not expand in number after bacterial infection. MAIT cells in the circulation are also reduced rather than expanded in patients with TB, sepsis and COVID-19 (10, 32, 35–38, 45, 46). MAIT cells may accumulate to slightly higher frequencies in tissues during some infections as has been shown in the BAL during tuberculosis or peritoneal cavity during spontaneous bacterial peritonitis (47, 48), but given the magnitude of changes reported this could be explained by recruitment from circulation rather than TCR-driven expansion. Based on the results presented here, it seems likely that MAIT cells do not mount large proliferative responses in humans and NHPs as part of their typical response to antigenic stimulation. In other words, our data raise the hypothesis that the lack of proliferation and downregulation of cytokine producing ability after strong antigenic challenge may not represent a defect but normal MAIT cell biology *in vivo* in humans and NHPs. That is not to diminish the importance of MAIT cells in host defense, as human MAIT cell deficiency has been shown to lead to increase susceptibility to viral and bacterial infections (49). We suggest the lack of MAIT cell responses reported here and in the above-mentioned literature is a beneficial adaptation of MAIT cells specifically in response to sudden and large increases in TCR ligand availability. Given that MAIT cells are likely persistently stimulated by antigens derived from the microbiota, there may be multiple checkpoints put on their ability to expand in number through large clonal bursts and produce inflammatory cytokines in order to prevent immunopathology.

It is not clear why the outcome of 5-OP-RU treatment of Mtb infection is so different in mice versus macaques. This may reflect a fundamental difference in MAIT cell biology between the two species. However, it could also be due to differences in MAIT cell extrinsic factors such as the microbiota composition or MAIT cell stimulation history. For example, at baseline murine MAIT cells in SPF mice are very low or negative for PD-1, while MAIT cells in macaques express significant levels of PD-1, indicating that there may be major differences in the levels of ambient TCR signaling between mice and NHPs. Therefore, mouse and macaque MAIT cells in the lungs may be in different functional states due to differences in the levels of persistent TCR stimulation at steady state. Additional studies are needed to better understand the mechanisms that regulate MAIT cell expansion in macaques so that strategies to induce large populations of MAIT cells *in vivo* can be developed, assuming a large number of activated MAIT cells is safe in NHPs and humans. We found that 5-OP-RU instillation led to a striking upregulation of Ki-67 in MAIT cells in both lung and blood, indicating that MAIT cells were likely induced to enter cell cycle. A reasonable hypothesis is that MAIT cell turnover was balanced by cell death, resulting in no net increase in cell number. It is possible that other adjuvants or different antigen delivery dynamics such as sustained low level 5-OP-RU stimulation may lead to MAIT cell expansion *in vivo.* Lastly, we have only used one MAIT antigen in this study, so it will be important to test other MR1 ligands for their ability to drive MAIT cell expansion in macaques.

Given the inability to expand MAIT cells, we could not test the hypothesis that large populations of MAIT cells can be protective against TB in this macaque study. Therefore, it remains possible that MAIT cell targeting therapies can be beneficial in Mtb infection, but this study highlights the significant barriers to exploring the role of MAIT cell-based therapeutics and vaccinations *in vivo.* Although the mouse model is a powerful tool that has revealed much about MAIT cell biology (50), there may be important instances where MAIT cell responses in mice may not be representative of responses in macaques and by extension also not representative of *in vivo* MAIT cell responses in humans. These data indicate it is important that hypotheses regarding MAIT cell biology that are developed in mice be further explored in humans if possible and NHP models when needed. The development of strategies to induce large populations of highly functional MAIT cells in macaques is needed to evaluate their clinical potential as targets of vaccines and therapeutics in humans.

## ACKNOWLEDGMENTS

We are grateful to Dr. Rashida Moore and Dr. Richard Herbert for providing veterinary care for animals in this study. We thank all staff members of the National Institutes of Allergy and Infectious Diseases (NIAID), Comparative Medicine Branch Animal Biosafety Level 3 facility for their technical support. This work was supported by the Intramural Research Program of the NIAID. G.J.F. is supported by P01AI056299 and R37AI112787. The members of the NIAID/DIR Tuberculosis Imaging Program are Janard L. Bleach, Ashley L. Butler, Emmuanual K. Dayao, Joel D. Fleegle, Felipe Gomez, Michaela K. Piazza, Katelyn M. Repoli, Becky Y. Slone, Michelle K. Sutphin, Alexandra M. Vatthauer, Laura E. Via, April M. Walker, Danielle M. Weiner, and Michael J. Woodcock.

## AUTHOR CONTRIBUTIONS

Conceptualization and Methodology, S.S. and D.L.B.; Investigation, S.S., N.E.L., K.D.K., D.E.D., S.N., F.G., J.D.F., NIAID/DIR TBIP and L.E.V.; Resources, S.O., C.S.L., A.S., G.J.F. and C.E.B.; Original Draft, S.S. and D.L.B.; Writing – Review & Editing, S.S., K.D.K., S.N., F.G., C.S.L., A.S., G.J.F., L.E.V., C.E.B. and D.L.B.; Visualization, S.S., S.N., F.G. and J.D.F.; Supervision, D.L.B., A.S., L.E.V. and C.E.B.; Funding Acquisition, D.L.B., A.S., G.J.F., L.E.V. and C.E.B.

## FIGURE LEGENDS

**Supplemental Figure 1. 5-OP-RU treatment does not cause substantial alterations in the intestinal microbiota.**

(**A**) Alpha diversity for each sample was estimated using the Shannon index. The left panel represents the diversity of each animal over the experimental timeline and PBS or 5-OP-RU treatment period is highlighted in grey. The right panel shows the pooled pre-infection (week −4/-2, 0), infected (week 4) and treated timepoints (week 8, 12 and necropsy) for the two groups. Statistical testing performed using Student’s t-test did not find any significant differences in alpha diversity between the timepoints for either group. Animals and groups can be identified as indicated in the key in B. (**B**) Beta-diversity analysis was performed using the Bray-Curtis dissimilarity matrix on the microbiota composition of the fecal samples collected following PBS or 5-OP-RU treatment. PERMANOVA was utilized to test for statistically significant differences in the microbiomes of the two treatments and was found to be not statistically significant (p-value = 0.055). (**C**) Relative abundance of phyla identified in the fecal samples of each study animal is represented. (**D**) LEfSe analyses depict bacterial families that are differentially abundant between the microbiota of pre-infection and treated timepoints in the PBS and 5-OP-RU groups and between the microbiomes of PBS and 5-OP-RU treated animals (LDA > 2, p < 0.05).

**Supplemental Figure 2. Association between the two methods used to estimate bronchoconstriction.**

Cross sections are the result of multiplying the diameter of the bronchus on coronal and sagittal views at a proximal point of the bronchus where constriction usually occurs. Volume was measured from the carinal bifurcation extending linearly 2.3 centimeters along the bronchus. This is the volume of the bronchus directly constricted by the enlargement of the peri-carinal lymph nodes.

## MATERIALS AND METHODS

### Rhesus macaques

Sixteen healthy rhesus macaques originally from the NIAID breeding colony on Morgan Island were selected for this study and were tuberculin skin test negative. Animals were housed in nonhuman primate biocontainment racks and maintained in accordance with the Animal Welfare Act, the Guide for the Care and Use of Laboratory Animals and all applicable regulations, standards, and policies in a fully AAALAC International accredited Animal Biosafety Level 3 vivarium. All procedures were performed utilizing appropriate anesthetics as listed in the NIAID DIR Animal Care and Use Committee (ACUC) approved animal study proposal LPD-25E. Euthanasia methods were consistent with the AVMA Guidelines on Euthanasia and endpoint criteria listed in the NIAID DIR ACUC approved animal study proposal LPD-25E.

### Mtb infection and 5-OP-RU treatment

Animals were infected with 120-150 colony forming units (CFU) of a *H37Rv* strain of Mtb. For infection, animals were anesthetized and 2 ml of PBS containing the bacteria were bronchoscopically-instilled into the right lower lung lobe. Infection dose was confirmed by plating of aliquots onto 7H11 agar plates. For the treatment during Mtb infection, 5 μg/kg of body weight of 5-OP-RU (14) diluted in 2 ml of PBS was administrated intratracheally once per week for 9 weeks beginning at 6 weeks post-infection. In PD-1 blockade experiment, uninfected animals were treated intratracheally with 0.5 μg/kg of body weight of 5-OP-RU and 100 μg of CpG ODN 2006 (InvivoGen) mixed with 10 mg/kg of body weight of either rhesus macaque IgG_4_ isotype control antibody (DSPR4) obtained from the NHP Reagent Resource or anti-PD-1 antibody (humanized clone EH12 kappa variable domains with rhesus macaque kappa and IgG_4_ constant regions) (51).

### PET/CT scanning and data analysis

Rhesus were imaged prior to infection and every two weeks beginning at 5 weeks post-infection for a maximum of 7 PET/CT scans (Figure 1A). The imaging studies were conducted with an optimized [^18^F]-FDG dose (0.5 mCi/kg) administered intravenously as previously described (52). A 360-projection CT scan of the lungs was acquired during a ~50 second breath hold on a LFER 150 PT/CT scanner (Mediso Inc, Budapest, Hungary). A 20-minute PET dataset/per field of view was acquired during mechanical ventilation and the raw CT and PET data were reconstructed using the Nucline software (Mediso, Inc, Budapest, Hungary) to create individual DICOM files that were co-registered using MIM Maestro (v. 6.2, MIM Software Inc, Cleveland, Ohio). A lung volume of interest (VOI) was defined on the CT image and the VOI was transferred to the PET image as previously described to determine the total [^18^F]-FDG uptake referred to as the lung total lesion glycolysis (TLG) (52). The analysis of disease burden included abnormal TLG of the lung, and a consistent region of the hylar and subcarinal LNs of each animal with standardized uptake value above background. The pulmonary LN [^18^F]-FDG uptake was measured by creating a VOI in MIM in the LN with uptake in the region surrounding the carina as described by White et al (53). Two readers independently performed image analysis for each animal. Threedimensional projections were generated using Osirix v 5.9 software (Pixmeo, Geneva, Switzerland).

Two approaches were taken for measuring the bronchial constriction observed in the infected animals. In the first, the diameters of the most constricted region of main right and left bronchi lumens were measured on the axial plane of CT scan with the embedded MIM ruler tool from anterior to posterior and from left to right. The measurements were compared to the baseline scan measurements at that axial plane to calculate a percentage of occlusion. The resulting percentages were averaged to get the mean percentage of occlusion at each time point. In the second method, the volume of the bronchus from the carinal bifurcation to 2.3 cm below from the carinal bifurcation (the area observed to contain the regions of occlusion in the macaques on study) was calculated by using the region grow feature of MIM set to capture the bronchi lumen (HU −1024 to −700) which was manually adjusted if small airway regions were missed by the program. The volumes of the right and left lumens were added together and compared to the baseline combined volume of the bronchi, and the percentage occlusion calculated. See also Supplemental Figure 2.

### Cell isolation and *in vitro* stimulation

Blood samples were collected in EDTA tubes and PBMCs were isolated by Ficoll-Paque (GE Life Sciences) density centrifugation. Broncho-alveolar lavage (BAL) samples were passed through a 100 μm cell strainer, pelleted, and counted for analysis. Lung robes and tracheal/bronchial tissues were minced using a GentleMACS Tissue Dissociator (Miltenyi Biotec) and were enzymatically digested in a shaker incubator at 37°C for 40 minutes in RPMI medium containing 1 mg/ml Collagenase D (Roche-Diagnostics), 1 mg/ml hyaluronidase and 50 U/ml DNase I (Sigma Aldrich). Suspensions were then passed through a 100 μm cell strainer and enriched for lymphocytes using a 40% Percoll density gradient centrifugation. Lymph nodes and spleens were dissociated using a GentleMACS Tissue Dissociator. Granulomas were individually resected from the lungs and samples used for flow cytometry analysis were pushed through a 100 μm cell strainer. Aliquots from all samples were serially diluted and plated on 7H11 agar plates for CFU quantification. Cells were stimulated in X-vivo 15 media (Lonza) supplemented with 10% FCS at 37 °C with either PMA/ionomycin (Leukocyte activation cocktail, BD Biosciences) for 3 hours, 5OP-RU (50 nM) or MHC-I (36) and MHC-II (MTB300) (54) Mtb peptide megapools (1 μg/ml and 2 μg/ml respectively) for 6 hours in the presence of brefeldin A and monensin (eBioscience). Cells were assessed for intracellular cytokine production as described below.

### Flow cytometry

Tetramer stains were performed by incubating 1×10^6^ cells at 37°C for 30 minutes with rhesus macaque MR1/5-OP-RU tetramer in X-vivo 15 media containing 10% FCS and monensin. Tetramers were produced by the NIAID tetramer core facility (Emory University, GA). Fluorochrome-labeled antibodies used for flow cytometric analysis are listed in Supplemental Table 1. Surface antigens and dead cells were stained in PBS + 1% FCS + 0.1% sodium azide for 20 minutes at 4 °C. For intracellular cytokine and transcription factor staining, cells were fixed and permeabilized with the Foxp3 Transcription Factor Staining Buffer Kit (eBioscience) and stained for 1 hour at 4 °C. Samples were acquired on a FACSymphony (BD Biosciences), and data were analyzed using FlowJo 10 (Treestar).

### Measurement of Mtb-specific IgG

Ninety-six-well ELISA plates were coated with Mtb whole cell lysate (strain H37Rv, BEI Resources) at 10 μg/ml diluted in PBS for 1 hour at 37°C. The plates were washed and blocked overnight at 4°C with block buffer (5% milk powder + 4% whey buffer in PBS Tween-20). Plates were then washed, and plasma samples were added at a serial 1:3 dilutions starting at a 1:10 dilution with 4% whey buffer and incubated for 1 hour at 37°C. After washing, plates were incubated with goat anti-monkey IgG (H+L)-HRP (Novus Bio) was added at 1:1000 dilution in 4% whey buffer for 1 hour at 37°C. Plates were washed and 1-Step Ultra-TMB ELISA Substrate Solution (Thermo Scientific) was added to develop the plates. The reaction was stopped by adding 0.5 M sulfuric acid, and the OD measured at 450 nm. The OD value of each pre-infection baseline was subtracted from each post-infection timepoint sample in order to calculate Mtb-specific IgG levels.

### Microbiota analyses

One pre-infection fecal sample was collected 2 to 4 weeks prior to infection and another on the day of infection. Additional samples were collected at weeks 4, 8, 12 post-infection and at necropsy. All samples were stored at −80 °C until completion of experiment. DNA was extracted from ~0.05g of fecal material using QIAamp Fast DNA stool Mini kit (Qiagen, Hilden, Germany) and the V4 region of the 16s rRNA gene was amplified with primers 5’-TCGTCGGCAGCGTCAGATGTGTATAAGAGACAGGTGCCAGCMGCCGCGGTAA-3’and 5’-GTCTCGTGGGCTCGGAGATGTGTATAAGAGACAGGGACTACHVGGGTWTCTAAT-3’ and sequenced as previously described (55). The raw reads were demultiplexed, denoised and filtered for chimeras using the DADA2/QIIME2 pipeline (version 2-2020.2) (56). The processed data resulted in an average of ~55,000 reads/sample. Alpha and beta-diversity analyses were performed using Shannon and Bray-Curtis dissimilarity indices respectively on read data rarefied to a depth of 40,000 reads/sample. Taxonomic classification was performed utilizing QIIME2 and the Silva database release 132 (57). Differentially abundant taxa were identified using Linear discriminant analysis (LefSe) and filtered for linear discriminant score (LDA) > 2 and p-value < 0.05 (58).

### Quantification and statistical analysis

All analyses were conducted using Prism 8 (GraphPad Software). Two-sample t test was used for two group comparisons and ANOVA was used for comparing multiple groups. Unless specifically denoted in the Figures, a p value < 0.05 was considered statistically significant. Data are presented as mean ± SEM.

## SUPPLEMTAL MATERIALS

**Supplemental Figure 1. 5-OP-RU treatment does not cause major alterations in the intestinal microbiota.**

**Supplemental Table 1. List of flow cytometry panels and antibodies used in this study.**

