## Supplemental Figure 1 for "Functional inactivation of pulmonary MAIT cells following 5-OP-RU treatment of non-human primates"

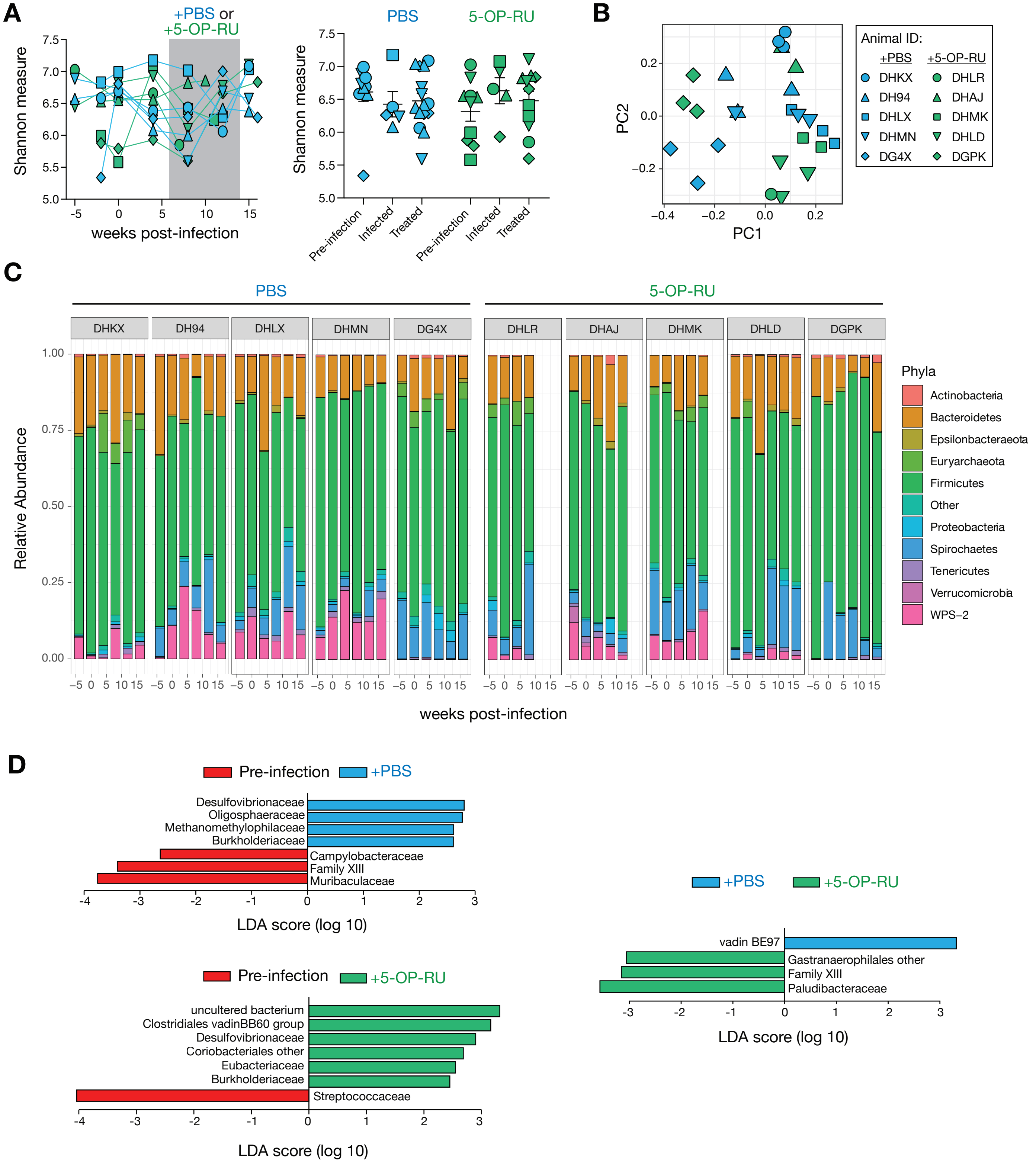


**Supplemental Figure 1. 5-OP-RU treatment does not cause major significant alterations in the intestinal microbiota.**
