## Supplemental Figure 2 for "Functional inactivation of pulmonary MAIT cells following 5-OP-RU treatment of non-human primates"

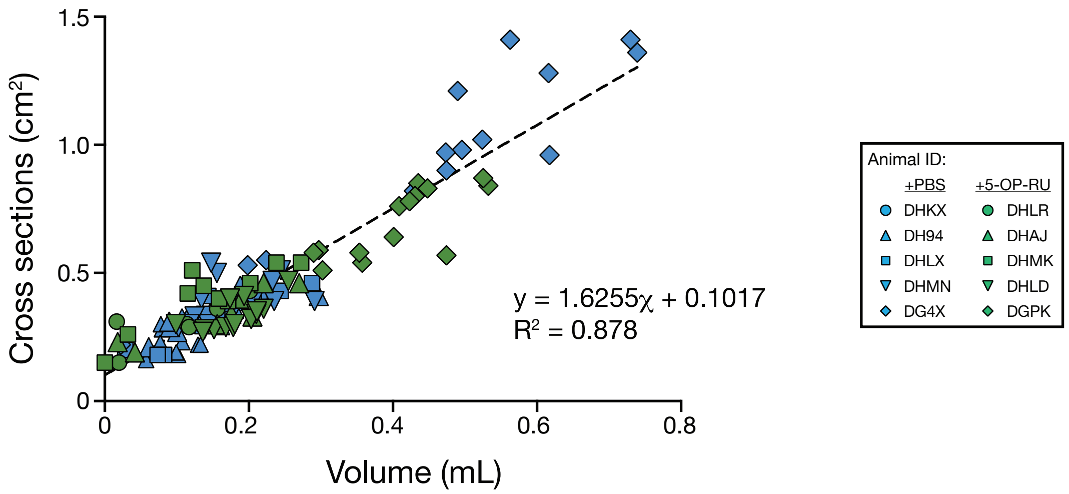


**Supplemental Figure 2. Association between the two methods used to estimate bronchial constriction.**
