## Supplemental Table 1 for "Functional inactivation of pulmonary MAIT cells following 5-OP-RU treatment of non-human primates"

**Supplemental Table 1. List of flow cytometry panels and antibodies used in this study.**

| | Antibody/Dye | Clone | Fluorochrome |
|---|---|---|---|
| **Panel 1** | CD8 | SK1 | BUV395 |
| | CD69 | FN50 | BUV496 |
| | CD3 | SP34-2 | BUV805 |
| | Ki-67 | B56 | BV421 |
| | CD4 | SK3 | BV510 |
| | Vα7.2 | 3C10 | BV711 |
| | PD-1 | EH12 | PE |
| | γδTCR | B1 | PE-Dazzle |
| | CD161 | HP3G10 | Pe/Cy7 |
| | MR1/5OP-RU tetramer | N/A | APC |
| | Live/Dead dye | N/A | Fixable Viability Dye 780 |
| **Panel 2** | IFN-γ | Mab11 | BUV395 |
| | CD4 | SK3 | BUV737 |
| | CD3 | SP34-2 | BUV805 |
| | GM-CSF | BVD2-21C11 | BV421 |
| | CD8 (ex) | SK1 | BV510 |
| | Vα7.2 | 3C10 | BV711 |
| | TNF-α | Mab11 | FITC |
| | IL-17A | eBIO64DEC17 | PE |
| | γδTCR | B1 | PE-Dazzle |
| | CD161 | HP3G10 | Pe/Cy7 |
| | MR1/5OP-RU tetramer | N/A | APC |
| | Live/Dead dye | N/A | Fixable Viability Dye 780 |
| **Panel 3** | TNF-α | Mab11 | BUV395 |
| | CD4 | SK3 | BUV496 |
| | CD95 | DX2 | BUV737 |
| | CD3 | SP34-2 | BUV805 |
| | CD28 | CD28.8 | BV421 |
| | CD8 | SK1 | BV510 |
| | IFN-γ | Mab11 | BV711 |
| | Foxp3 | 150D | FITC |
| | PD-1 | EH12 | PE |
| | Live/Dead dye | N/A | Fixable Viability Dye 780 |
